# Benzamidine-Mediated Inhibition of Human Lysozyme Aggregation: Differential Ligand Binding in Homologous Proteins

**DOI:** 10.1101/2024.10.17.618876

**Authors:** Sree Hima, Chandran Remya, J Abhithaj, K.G. Arun, A. Sabu, D. M. Vasudevan, K.V. Dileep

**Affiliations:** Amrita Vishwa Vidyapeetham, Kochi, Kerala, India; Laboratory for Computational and Structural Biology, Jubilee Centre for Medical Research, Jubilee Mission Medical College and Research Institute, Thrissur, Kerala, India; Inter University Centre for Bioscience and Department of Biotechnology & Microbiology, Dr Janaki Ammal Campus Thalassery, Kannur University, Palayad 670661, India

**Keywords:** Human lysozyme, Benzamidine, Aggregation, Congo red, ANS, Thioflavin, Docking

## Abstract

Amyloid fibril formation is a hallmark of several protein misfolding diseases, including systemic hereditary amyloidosis (SHA), in which lysozyme aggregates into plaques, causing inflammation in various tissues. SHA is a rare disease with no current drug treatment options. In our efforts to identify potential therapeutics for SHA, we investigated the inhibitory effects of benzamidine (BEN) on the fibrillation of human lysozyme (HL). Multiple biophysical assays demonstrated BEN’s ability to effectively prevent amyloid formation. Intrinsic fluorescence measurements highlighted BEN’s interaction with HL. We inferred the binding mode of BEN to HL through ITC experiments, molecular docking, and molecular dynamics simulations, confirmed BEN’s binding at the active site, particularly near stretch-2 (residues 52-64), a key region in its anti-amyloidogenic activity. This interaction differed from the previously reported interaction with HEWL. Further, microscopy analyses, including scanning electron microscopy (SEM) and transmission electron microscopy (TEM), further supported these findings by showing reduced fibril formation and alterations in fibril morphology in the presence of BEN. Importantly, BEN exhibited no cytotoxic effects in HEK-293 cells, reinforcing its potential as a therapeutic candidate for amyloidosis. These results provide strong evidence of BEN’s anti-amyloidogenic activity and offer a foundation for future drug development targeting lysozyme amyloidosis.

## Introduction

Amyloidosis encompasses a diverse spectrum of proteinopathies characterized by the extracellular deposition of insoluble amyloid fibrils. These fibrils are formed due to the abnormal misfolding of proteins when they lose their native structure and function. They are often linked to cellular toxicity, leading to progressive tissue and organ dysfunction **[1]**. Based on the distribution of amyloid deposits, they are classified into systemic and localized **[2, 3]**. Formation and deposition of amyloid fibrils is a hallmark for many diseases like Alzheimer’s disease (AD), Parkinson’s disease (PD), creutzfeldt-Jakob disease (CJD), systemic hereditary amyloidosis (SHA) etc. Inhibiting the formation of amyloid fibrils has become one of the important strategies to treat amyloid-related diseases.

Lysozyme amyloidosis is a type of systemic amyloidosis characterized by the aggregation of lysozyme protein within the body, resulting in multiorgan dysfunction. It is a hereditary autosomal dominant disease marked by the accumulation of human lysozyme (HL), a 14 kDa bacteriolytic enzyme, distributed in various organs such as the gastrointestinal tract, lymph nodes, spleen, thyroid, kidney, liver, lungs, and salivary glands **[4]**. The disease manifests through diverse pathological symptoms, including hepatic rupture, renal failure, lymphadenopathy, sicca syndrome, purpura, gastrointestinal issues, and petechiae **[5, 6]**. Patients with lysozyme amyloidosis often exhibit massive lysozyme deposits in the kidney interstitium, perivascular areas, and glomeruli **[7]**. Currently, there is no established cure for lysozyme amyloidosis, and the available treatment only focuses on alleviating symptoms to improve patients’ quality of life. Therefore, the development of effective therapeutic strategies aimed at either blockade of amyloid fibrils or clearance of amyloid aggregates are highly demanding.

Benzamidine (BEN), a reversible inhibitor of trypsin, trypsin-like enzymes, and serine proteases is widely used not only in the biochemical research but also in the painful and inflammatory conditions of the oral cavity, such as infections and gingivitis **[8]**. The crystal structure of BEN in complex with hen egg white lysozyme (HEWL) has already been reported **[9]**. Given the stable binding of BEN demonstrated through X-ray crystallography, we hypothesized that BEN could similarly bind to HL, as both of the proteins shares high structural similarities. We also hypothesized that BEN’s interaction with HL would disrupt the aggregation process, preventing the misfolding events. To test our hypothesis, we performed series of biophysical and molecular modeling studies.

## Materials and methods

### Protein and chemicals

All chemicals used in this study were of analytical reagent grade and used as received without additional purification. The HL, BEN and Congo Red (CR) were purchased from Sigma-Aldrich, India, Thioflavin T (ThT) and 8-Anilinonaphthalene-1-sulfonic acid (ANS) were obtained from Tokyo Chemical Industry (Japan).

### Preparation of HL aggregates

Stock of HL was prepared by dissolving the protein in citrate-phosphate buffer (0.1 M, pH 2.2; hereinafter referred to as CA-PB buffer). To induce the amyloid formation, the HL solution was incubated for 120 h at 60°C in a shaking incubator (250 rpm). To check the action of BEN on HL fibrils, the same experiment was repeated in presence of different concentrations of BEN (25, 50, 100, 200, 400 and 800 μM), prepared in double distilled water (ddH_2_O)).

### ThT fluorescence assays

The formation of HL fibril was monitored by a characteristic increase in ThT fluorescence intensity. After 5 days of incubation, ThT (dissolved in ddH_2_O) (final concentration of 20 μM) was added to the HL samples (100 μM) in presence and absences of different concentrations of BEN. The fluorescence intensity of HL fibril was measured by multimode plate reader (Tecan Infinite F500) with parameters as follows: the excitation wavelength was 440 nm; the emission spectrum was recorded at 460 and 600 nm.

To clarify the quenching mechanism, both the Stern-Volmer quenching constant (K_SV_) and the quenching rate constant (Kq) were determined using the Stern-Volmer equation.

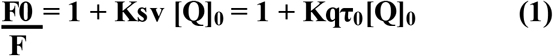

where F0 and F represent the fluorescence intensities of HL in the absence and presence of BEN, respectively. [Q] is the concentration of BEN, and τ_0_ is the fluorescence lifetime of the protein, typically valued at 10^−8^ seconds **[10]**.

### Congo Red (CR) binding assay

The fibrillation of HL was evaluated using a CR binding assay **[11]**. A 20 μM CR working solution was prepared by diluting CR in a 20 mM sodium phosphate buffer containing 50 mM NaCl (pH 7.4). This solution was then mixed with the protein samples (HL incubated at 60°C for 120 hours) at a 4:1 molar ratio, both in the absence and presence of BEN, and incubated for 30 minutes in the dark. The absorption spectra of each sample were measured between 400 and 700 nm using a multimode plate reader (Tecan Infinite F500).

### ANS fluorescence assays

A 10 mM stock solution of ANS was prepared by dissolving ANS in ddH_2_O and filtered using Whatman filter paper and stored at 4°C for future use. Protein fibril solution (5 µM), both with and without BEN, were incubated with ANS at a molar ratio of 1:10 (protein to ANS ratio) for 15 minutes in the dark. The samples were then excited at a wavelength of 380 nm, and the emission spectra were recorded from 400 to 600 nm using multimode plate reader (Tecan Infinite F500). This setup allowed for the precise observation of the protein-ANS interaction and provided insights into the binding dynamics in the presence and absence of BEN.

### Intrinsic Fluorescence Measurements

Intrinsic or tryptophan fluorescence intensities of the untreated HL and BEN with different molar ratios were monitored on the spectrofluorometer. The fluorescence spectra ranging from 305 to 400 nm were recorded upon exciting the samples at 270 nm (for tryptophan fluorescence). Both excitation and emission slits were set at 10 nm.

### Isothermal Titration Calorimetry

Thermodynamic evaluation of enzyme-ligand binding was conducted using the MicroCal PEAQ-ITC (Malvern) instrument at a controlled temperature of 25°C. For the experiment, 300 microliters of the protein solution (50 μM) were carefully loaded into the sample cell, while the syringe was filled with 60 microliters of the BEN solution (500 μM). To ensure accurate measurements, the ligand solution was degassed prior to the experiment to remove any air bubbles, which could interfere with the sensitive calorimetric measurements. The degassed BEN solution was gradually titrated into the sample cell. The first injection was set to a small volume of 0.4 μL to minimize initial disturbances, followed by subsequent injections of 3 μL each. A total of 13 injections were performed with a time interval of 150 seconds between each, allowing for proper equilibration and measurement of the heat change associated with each injection. The reference power was calibrated to 10 μcal, and the stirring speed was maintained at 750 rpm to ensure uniform mixing of the protein-ligand complex within the cell. After the experiment, the data acquisition and analysis were performed using MicroCal PEAQ-ITC Analysis Software, which enabled the calculation of key thermodynamic parameters, including the binding constant (K), enthalpy change (ΔH), entropy change (ΔS), and Gibbs free energy (ΔG).

### Molecular modeling studies

To investigate the binding interaction of BEN with HL, molecular docking was carried out using Schrödinger Maestro. The HL crystal structure (PDB ID: 1JSF) was selected for molecular docking studies. Prior to molecular docking, the protein was prepared by deleting the water molecules, adding polar hydrogens to the protein structure and minimizing energy by applying a cut of 0.30 Å using a OPLS 2001 force field. All the protein preparations were done using Protein Preparation Wizard module. The ligand BEN was prepared using LigPrep module. Before performing docking, two grids were generated on the protein to define the binding site. Using these grid information, two separate docking simulations were conducted. The first grid was set up based on the prior knowledge of how BEN binds to HEWL, we utilized this information to define the grid on HL. The grid was configured with a box dimension of 10 x 10 x 10 Å, centered on the binding pose of BEN.

The best energy poses from two different docking were selected for the MD simulations. The simulations were performed for 250 ns to assess the free energy of binding and residence of time of the ligand (RMSD). All MD simulations were performed by GPU accelerated Desmond software. Prior to MD simulations, the system was set up using the ‘System Builder’ in Desmond. The protein-ligand complex was immersed into an orthorhombic box with dimensions of 10 x 10 x 10 Å containing TIP3P explicit solvent model. The systems were further neutralized by adding Na + or Cl - ions and MD simulations were performed at a constant temperature of 300 K. Default settings were used for all other parameters.

### Electron Microscopic studies

The HL fibrils (100 μM), both in the presence and absence of BEN (800 μM), were prepared for electron microscopy analysis to study their morphological changes. For scanning electron microscopy (SEM), approximately 10 μL of the sample was deposited onto a freshly cleaved gold-palladium surface. The sample was then left to dry for 1 hour under ambient conditions. Morphological alterations of HL fibrils, with and without BEN, were examined using a TESCAN VEGA SEM. The surface topography and fibril network were carefully analyzed to assess any structural differences induced by BEN.

Transmission electron microscopy (TEM) was also employed to further investigate the ultrastructural changes. TEM images of the HL fibrils, with and without BEN, were captured using a JEOL JEM-1200EX high-resolution TEM (Japan). The samples were carefully cast onto copper grids and allowed to dry for 1 hour before imaging. High-resolution images were obtained, and both the length and diameter of the fibrils, in the presence and absence of BEN, were measured using Image-J software **[12]**. This allowed for a detailed comparison of fibril dimensions and aggregation patterns, providing insights into the potential inhibitory effect of BEN on HL fibril formation.

### Cytotoxicity assessment of BEN

The cytotoxicity of BEN was evaluated using the MTT assay on HEK-293 cells, which measures cell viability by detecting the enzymatic conversion of MTT (measured at 570 nm), a pale-yellow tetrazolium compound, into dark purple formazan crystals by mitochondrial succinate dehydrogenase. The HEK-293 cells were cultured in DMEM supplemented with 10% fetal bovine serum (FBS), 10,000 units/mL penicillin, 10 mg/mL streptomycin, and 25 mg/mL amphotericin-B in T-25 flasks. Cells were then trypsinized and seeded into 96-well plates at a density of approximately 20,000 cells per well. After 24 hours, the medium was replaced with fresh medium containing various concentrations of BEN (50, 100, 200, 400, and 800 μM), all dissolved in DMSO, ensuring a final DMSO concentration of 0.08%. Following 24-hour incubation, the MTT solution (5mg/mL) was added to each well, and the cells were incubated for an additional 2 hours to allow for formazan crystal formation. Later 100 µL of DMSO solution was added to dissolve the formazan crystals and the absorbance at 570 nm was measured using a multimode plate reader (Tecan Infinite F500). The cell viability was calculated as a percentage relative to the control group, which consisted of cells treated with 0.08% DMSO.

## Results and discussion

### Biophysical Insights Highlight BEN’s Inhibitory Effect on HL Fibrillation

The change in the fluorescence intensity of ThT served as a key marker to detect the fibril formation **[13]**. The changes in ThT fluorescence intensity of HL fibrils in the presence and absence of BEN suggests that the BEN could effectively block the fibril formation **(Figure 2A)** in a dose-dependent manner. At higher concentrations of BEN, the fibrillation effect of HL has been decreased significantly **(Figure 2B)**. Further the Ksv and Kq values for the interaction between BEN and HL were determined by fitting the data to the Stern-Volmer equation **(Figure 2C)**. The calculated Kq value provided insight into whether the quenching mechanism was static or dynamic. The quenching constant Ksv was 2.4 × 10^3^ L mol^−1^, and the quenching rate constant Kq was 1.53 × 10^11^ L mol^−1^ s^−1^. Since the Kq value exceeds the diffusion-limited rate constant for biomolecules (2 × 10^10^ L mol^−1^ s^−1^), it suggests that the quenching mechanism is static **[14]**, attributed to the formation of a strong complex between BEN and HL.

**Figure 1.**
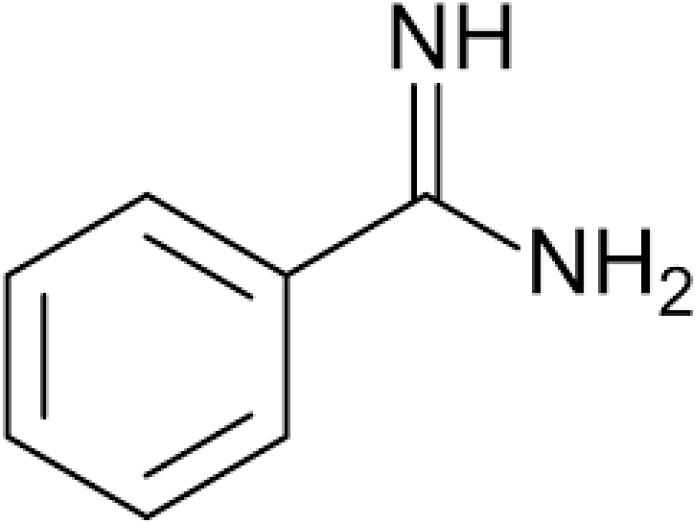
Chemical structure of BEN.

**Figure 2:**
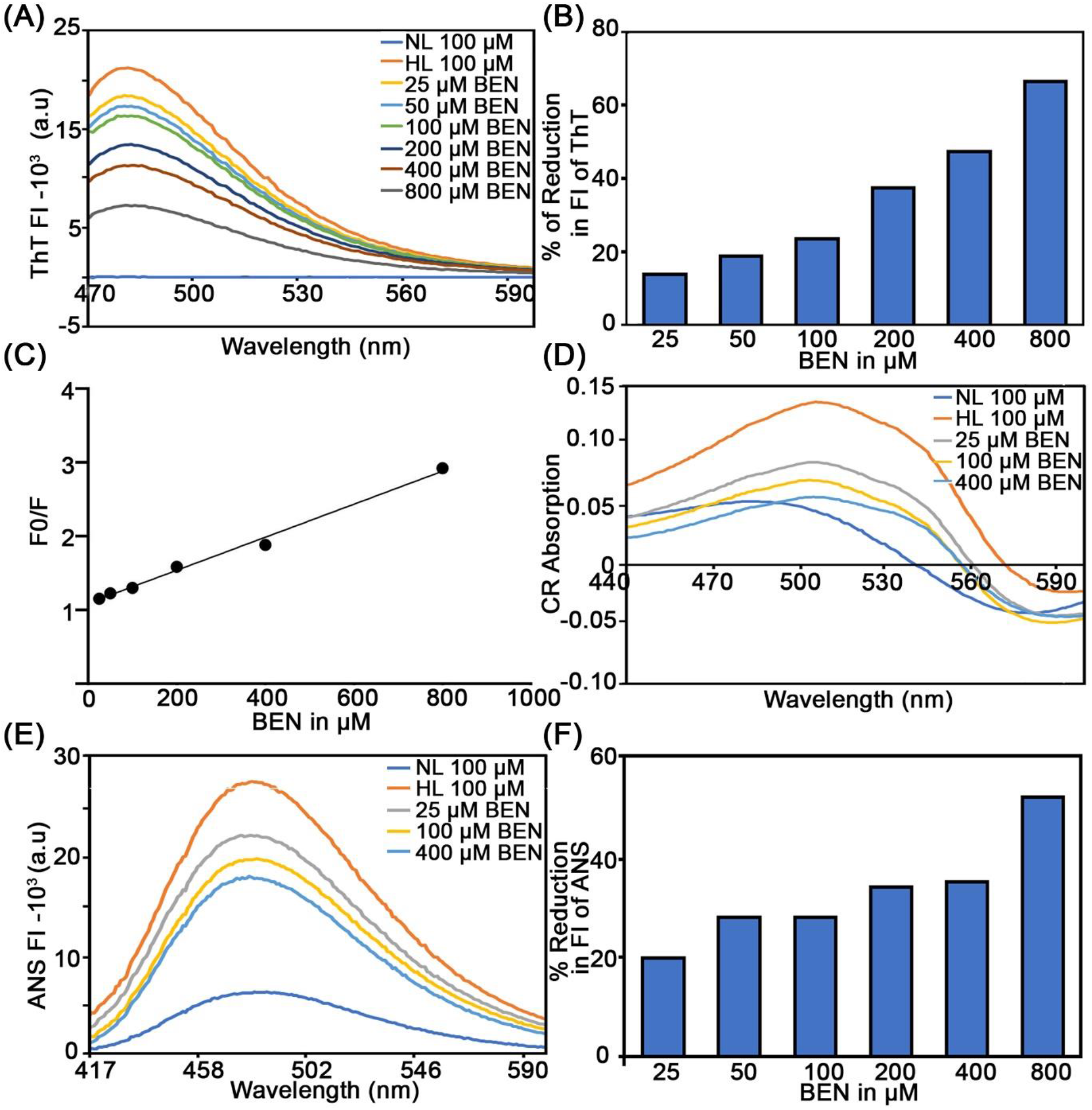
Inhibition of HL fibrillation by BEN as analyzed through various biophysical assays.**(A)**ThT fluorescence spectra showing the aggregation of HL in the absence and presence of increasing concentrations of BEN (25–800 μM).**(B)** Quantitative analysis of the % of reduction in ThT fluorescence intensities at 483 nm, indicating the inhibition of HL fibril formation by BEN in a dose-dependent manner.**(C)** Stern-Volmer plot depicting the fluorescence quenching of HL by BEN.**(D)** CR absorption spectra of HL fibrils in the absence and presence of BEN.**(E)** ANS fluorescence spectra of HL fibrils in the absence and presence of BEN, showing the effect of BEN on the hydrophobic exposure of HL fibrils.**(F)** Quantitative representation of the reduction in ANS fluorescence intensities at 480 nm. NL and HL indicate native and human lysozyme respectively.

Similar to the ThT molecule, CR also binds specifically to the flat β-sheet structures formed as a part of fibril formation **[15]**. The interaction between CR and the flat β-sheet causing a increase as well as a red shift in the maximum absorbance **[16]** suggesting the formation of fibrils compared to native lysozyme. The CR absorption spectra exhibited a red shift to 545 nm in HL fibrils compared to native lysozyme. In the presence of BEN, the absorbance values decreased in a dose-dependent manner **(Figure 2D)**. These results indicate that BEN disrupts the formation of cross β-sheet structures in HL protein, thereby inhibiting fibril formation.

The ANS fluorescence assay can detect the early stages of fibril formation by binding to the hydrophobic regions on macromolecules **[17-20]**. A blue shift in the ANS emission indicates high-affinity binding to solvent-exposed hydrophobic patches within amyloid fibrils **[21]**. In the current study, when ANS was treated with samples such as HL, both in the presence and absence of BEN, a blue shift in ANS emission was observed **(Figure 2E)**. Each sample exhibited maximum fluorescence intensity at 480 nm, indicating that the hydrophobic surface groups of HL were fully exposed during fibrillation. As BEN concentration increased, a concentration-dependent decrease in ANS fluorescence intensity was noted **(Figure 2F)**. This suggests that BEN effectively inhibits HL amyloid fibrillation, by arresting the exposure of hydrophobic patches to the solvent, a step in the HL aggregation process.

### Evidences suggests that BEN preferentially binds to the HL’s active site, halting fibrillation

The intrinsic fluorescence of the protein can be used to evaluate the structural, physicochemical, and functional properties **[22]**. HL contains five Trp (W) residues at the positions of 28, 34, 64, 109, and 112. Of these residues, W34 is located on the surface of HL, while W64 and W109 are positioned near the active site cleft. In contrast, W28 and W112 are buried within the protein structure, making them inaccessible to protein fluorophores or quenchers. Variation in the microenvironment of tryptophan residues and monitoring the intrinsic fluorescence of a protein allows assessment of its structural characteristics **[23, 24]**. Additionally, probing the accessibility of protein fluorophores to quenchers provides insights into binding interactions. In this study, HL was titrated with increasing concentrations of BEN in a distilled water at 298 K **(Figure 3A)**. It was observed that with the addition of BEN, the fluorescence intensity of HL gradually decreased, indicating that the interactions of BEN with HL caused quenching in the fluorescence. Based on the positions of the Trp residues and the observed reduction in quenching in the presence of BEN, we hypothesized that BEN may bind near to W34 or within the active site. In a previously reported crystallographic structure of HEWL complexed with BEN, the binding site of BEN was found near F34 (in HL, F34 is replaced by W34). Additionally, ligands such as gallic acid, , benzoic acid, pyrogallol **[25]**, vanillin **[15]** and Eugenol **[26]** have been reported to bind to the active site of HEWL. Considering the structural similarities between these ligands and BEN, it is reasonable to assume that BEN may also bind to the active site, similar to these molecules. Intrinsic fluorescence measurements suggested the possibility of multi-site binding for BEN. To confirm this, we conducted ITC experiment.

**Figure 3:**
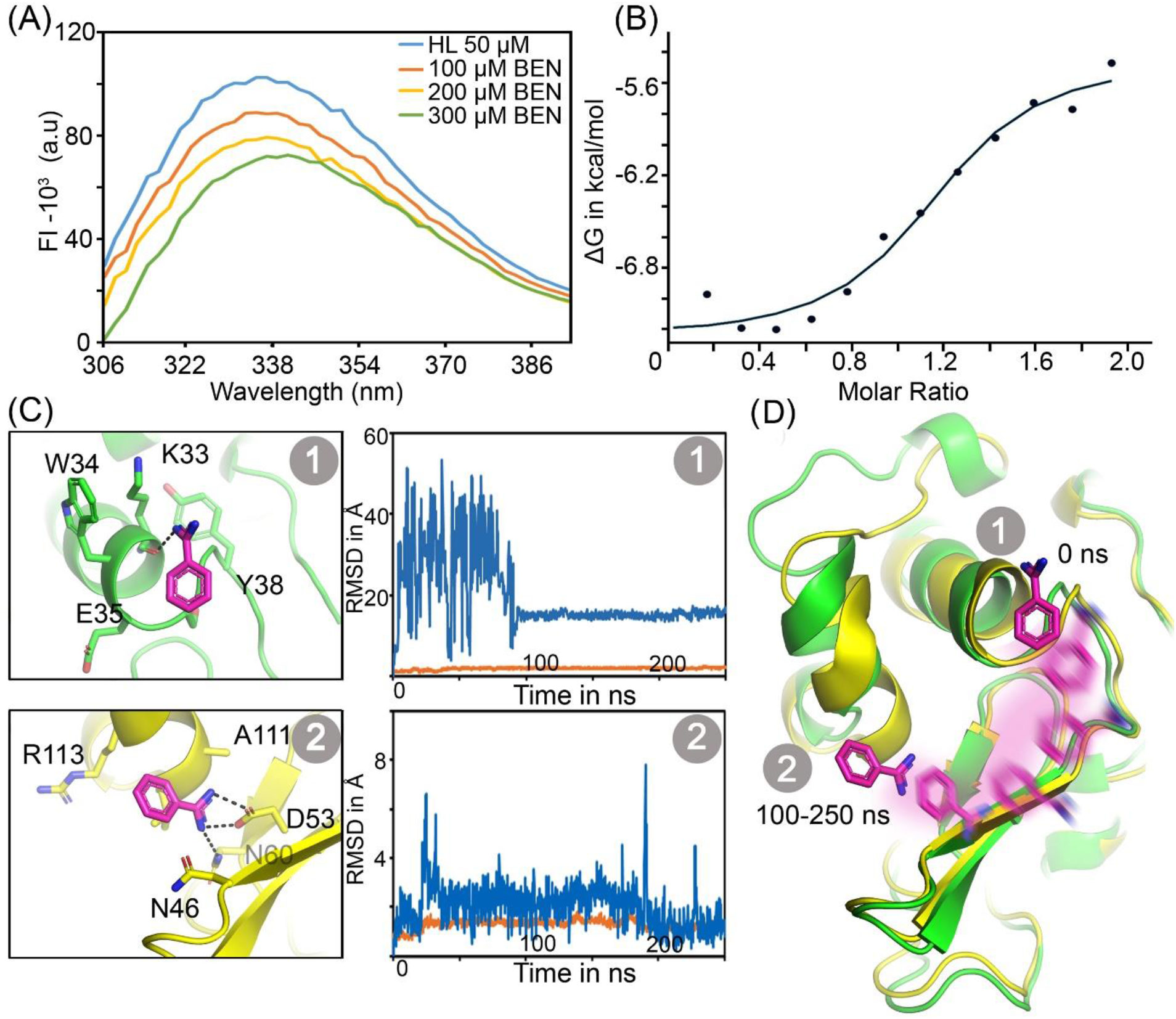
Binding mechanism of BEN with HL. **(A)** Intrinsic fluorescence emission spectra of lysozyme with varying concentrations of BEN (100, 200, and 300 μM). **(B)**The thermodynamic profile of BEN-HL interaction derived through ITC. **(C)** Binding modes of BEN at two distinct sites (represented as 1 and 2) on HL. The left-top panel shows the binding mode predicted through molecular docking, while the left-bottom panel illustrates the self-explored binding mode identified during MD simulations. In both cases, BEN is shown as pink sticks, and the proteins are represented by green and yellow sticks and cartoons. The corresponding RMSD values of the protein (orange lines) and ligand (blue lines) interactions are depicted in the right-top and bottom panels. **(D)** The binding mode of BEN and its self-exploratory binding trajectory at the active site during a 250 ns simulation are also shown.

Our ITC experiment deduced the thermodynamic parameters governing the binding between HL and BEN. The ITC results demonstrated that BEN binds strongly to HL, with a binding constant (K) of 6.47×10^5^ M^−1^**(Figure 3B and S1)**. The stoichiometry of binding (N) was determined to be 1.14, indicating nearly one BEN molecule interacts with each HL molecule. The thermodynamic parameters indicate an energetically favorable binding process, with a negative Gibbs free energy (ΔG = -7.67 kcal/mol), which reflects a spontaneous interaction. The exothermic nature of the interaction is highlighted by the negative enthalpy change (ΔH = -1.82 kcal/mol), suggesting the formation of favorable interactions, such as hydrogen bonds or van der Waals forces, between BEN and HL during complex formation. The entropy change deduced from the ITC experiment is ΔS = -5.85 cal/mol/deg **(Table 1)**.

**Table 1:** Thermodynamic parameters calculated for the binding of BEN from ITC analysis.

| Compound | N | K mol<sup>-1</sup> | T (K) | $\Delta S$<br>cal/mol/deg | $\Delta H$<br>cal/mol | $\Delta G$ in<br>kcal/mol |
| --- | --- | --- | --- | --- | --- | --- |
| BEN | 1.14 | 6.47 x 10<sup>5</sup> | 298.15 | -5.85 | -1.82 | -7.67 |

Both intrinsic fluorescence and ITC experiments not only confirmed the stable binding of BEN to HL but also indicated a single binding site. To identify the possible binding site, molecular docking studies were performed using the crystal structure of HL (PDB ID: 1JSF). Comparative insights from BEN’s known binding interactions with HEWL (PDB ID: 4NX6) helped us to pinpoint the binding region on HL. Docking results revealed that BEN binds to HL in a similar manner to its binding in HEWL, with an RMSD ≤ 1 Å between the ligands in the HEWL-BEN complex (4XN6) and the HL-BEN complex. The binding free energy of BEN to HL (−14.38 kcal/mol) was slightly weaker than its binding to HEWL (−19.21 kcal/mol). Structural analysis showed three hydrogen bonds in the HEWL-BEN complex, in which K33 and N37 forming direct hydrogen bonds with BEN. Residue F34 has been involved in a bridged hydrogen bond through a water molecule. However, in the HL-BEN interaction (as per our molecular docking results), some of these hydrogen bonds were absent **(Figure 3C)**. Notably, N37 in HEWL is replaced by G37 in HL, resulting in the loss of the side chain hydrogen bond with BEN. This difference in interactions likely accounts for the decrease in the binding free energies. In both proteins, hydrophobic and van der Waals interactions play a major role in stabilizing BEN in their binding sites.

Since all docking studies were performed using a rigid docking mode, we evaluated the stability of BEN (residence time) in the HL and HEWL complex through MD simulations. Over the course of 250 ns, the BEN-HEWL complex remained stable **(Figure S2)**, supported by three strong hydrogen bonds. However, in the BEN-HL complex, BEN initially exhibited large fluctuations and migrated from its original binding position, started exploring different regions on HL. After 100 ns, BEN identified the active site and remained stable for the rest of the simulation, forming key interactions with residues D53 and N60. Further to reconfirm the stability of BEN in the active site, we repeated the simulation without changing the parameters using a representative snapshot from the 100–250 ns range. This simulation further confirmed that BEN stayed within the HL active site, maintaining stable interactions with D53 and N60 throughout **(Figure 3C and D)**.

Previous studies have identified that the residues 22-47 (stretch-1), 52-64 (stretch-2), 85-90 (stretch-3), and 105-113 (stretch-4) are belongs to the aggregation-prone regions of HL**[27]**. The binding of BEN near to stretch-2 might be contributing to its anti-amyloidogenic activity. In conclusion, interaction of BEN to HL through hydrogen bonding, maintains conformational stability, and inhibits its aggregation, providing a potential mechanism for its anti-amyloidogenic effects.

### Microscopy Imaging suggested the effect of BEN on the fibrillation

Biophysical studies confirmed that BEN significantly reduces fibril formation. To further visualize the morphology of HL fibrils in the presence and absence of BEN, SEM and TEM analyses were performed. As expected, BEN at 800 µM significantly reduced fibril formation. The SEM images of HL fibrils in the absence of BEN appeared to large, continuous, and densely packed thread-like structures, indicative of successful and robust amyloid assembly. In contrast, the sample treated with BEN showed a notable reduction in fibril density and increased fragmentation **(Figure 4A)**. The fibrils in the presence of BEN appeared shorter and irregular, suggesting that BEN disrupts fibril growth and destabilizes fibril formation **(Figure 4B)**. This altered morphology confirms that BEN effectively inhibits the formation of HL amyloid fibrils.

**Figure 4:**
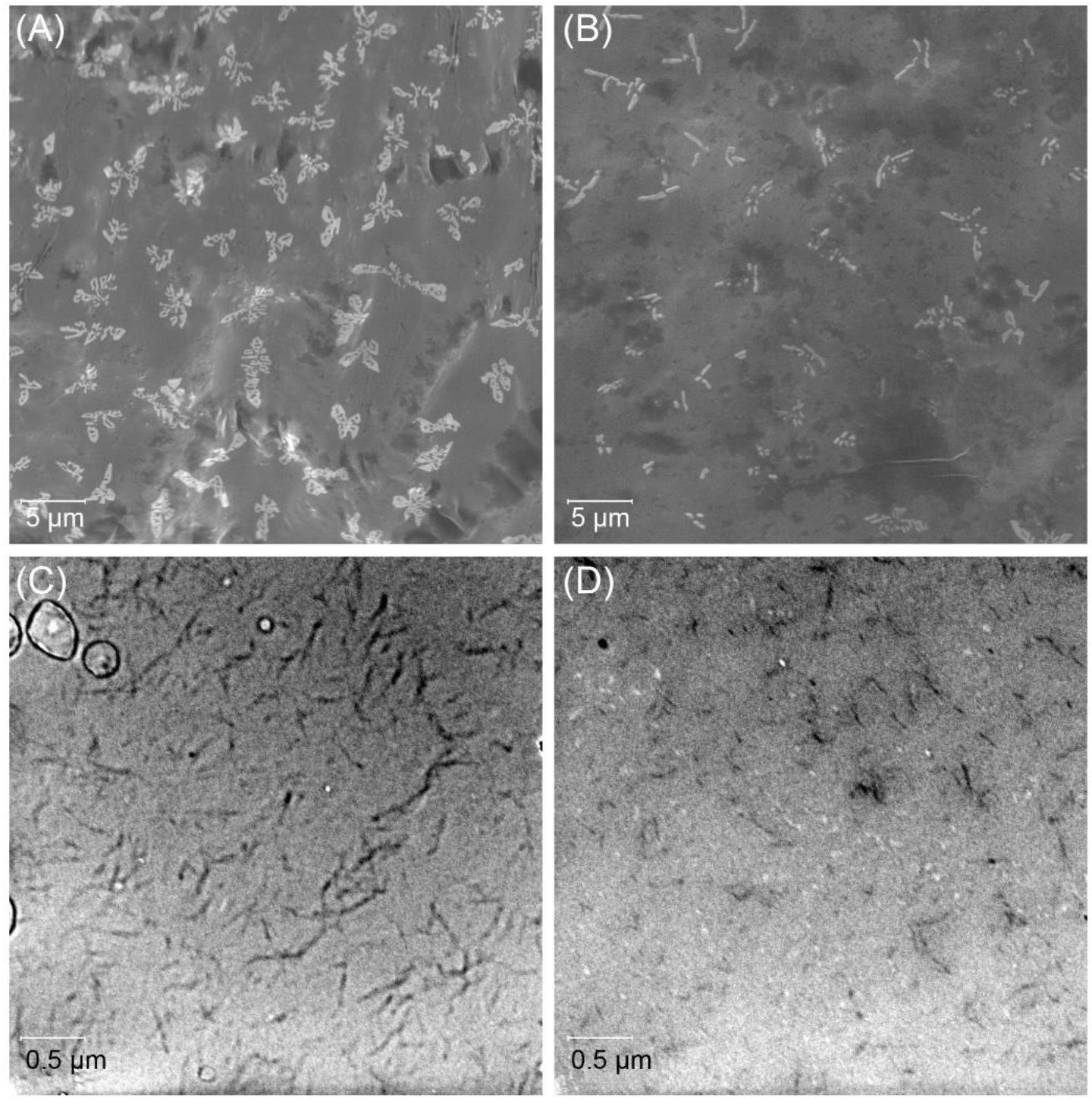
Effect of BEN on HL fibrillation as observed through microscopy imaging. SEM analysis reveals the morphological differences in HL fibrils **(A)** in the absence and **(B)** in the presence of 800 μM BEN. The TEM images further illustrate these differences, showing HL fibrils **(C)** without BEN and **(D)** with 800 μM BEN treatment. The scale bars are also indicated in each panel.

In the case of TEM, the presence of BEN (800 μM) in the HL provided significantly less dense, shorter and thinner fibrils **(Figure 4C)**. The reduction in both the number and length of fibrils (35-42µm) suggests that BEN interferes with the nucleation and elongation phases of fibril formation **(Figure 4D)**. The results intuitively confirmed that BEN could effectively inhibit the fibrillation process of HL.

### BEN is not cytotoxic in the HEK cell lines

Lysozyme amyloidosis is a disease primarily affecting the liver, kidney, and gastrointestinal tract **[5]**. To assess the cytotoxic nature of BEN, HEK-293 cells were incubated with varying concentrations of BEN for 24 hrs. The MTT assay performed on HEK-293 suggested that BEN even at 800 μM is not cytotoxic **(Figure 5)**, making it a potentially therapeutic candidate for inhibiting lysozyme aggregation. These findings strongly suggest that BEN as such or analogues of BEN could be explored further in amyloidosis treatment without adversely affecting kidney cells.

**Figure 5:**
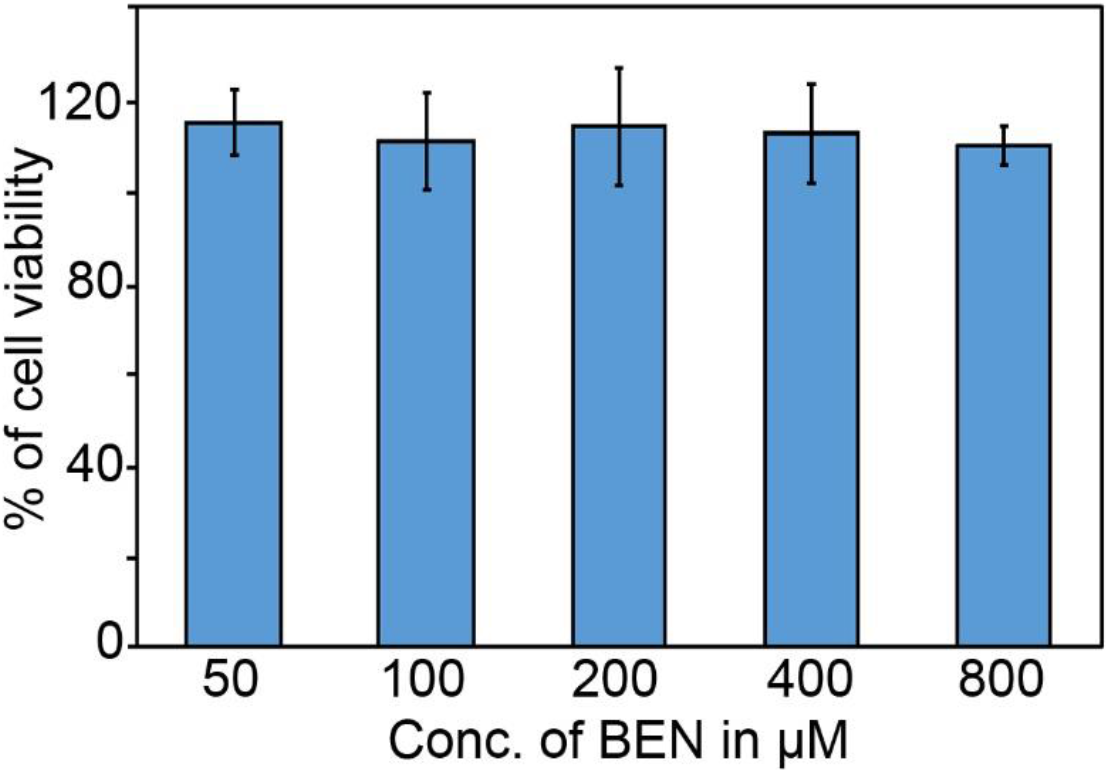
MTT assay for testing the cell cytotoxicity of BEN on HEK cell lines at different concentrations of BEN, *n=3*.

## Conclusion

In this study, we present comprehensive evidence of BEN’s inhibitory effect on HL fibrillation. A series of biophysical assays, including ThT, CR, and ANS fluorescence, revealed that BEN significantly disrupts fibril formation in a dose-dependent manner. Intrinsic fluorescence and ITC experiments confirmed BEN’s binding to HL, characterized by favourable thermodynamic parameters and a single binding site. Molecular docking and MD simulations further supported BEN’s interaction with key residues in the active site, particularly in the aggregation-prone stretch-2 region, suggesting a plausible mechanism for its anti-amyloidogenic activity.

Microscopy analyses (SEM and TEM) showed structural alterations in HL fibrils upon BEN treatment, with reduced fibril density and length, indicating disrupted nucleation and elongation. Moreover, BEN exhibited no cytotoxicity in HEK-293 cells at concentrations up to 800 μM, underscoring its potential as a therapeutic agent for lysozyme amyloidosis. These findings suggests that BEN could be a promising lead compound for the development of novel agents to prevent lysozyme aggregation, with potential applications in amyloidosis therapy.

## Acknowledgment

R.C. gratefully acknowledges the Chief Ministers Nava Kerala Postdoctoral Fellowship.

