## Supplementary material for "Benzamidine-Mediated Inhibition of Human Lysozyme Aggregation: Differential Ligand Binding in Homologous Proteins": Supplimentary Information

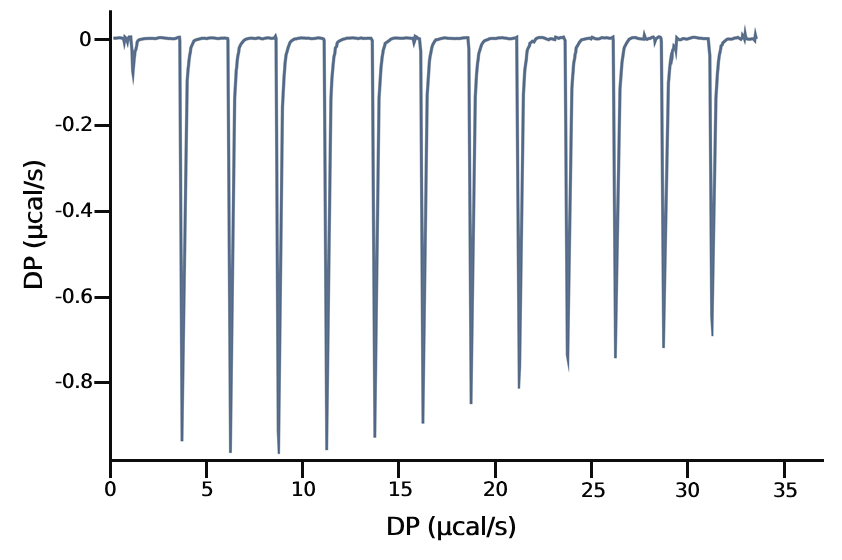

**Figure S1:** ITC analysis of HL with BEN. The exothermic peaks (Raw thermal power signal) obtained during the experiment.

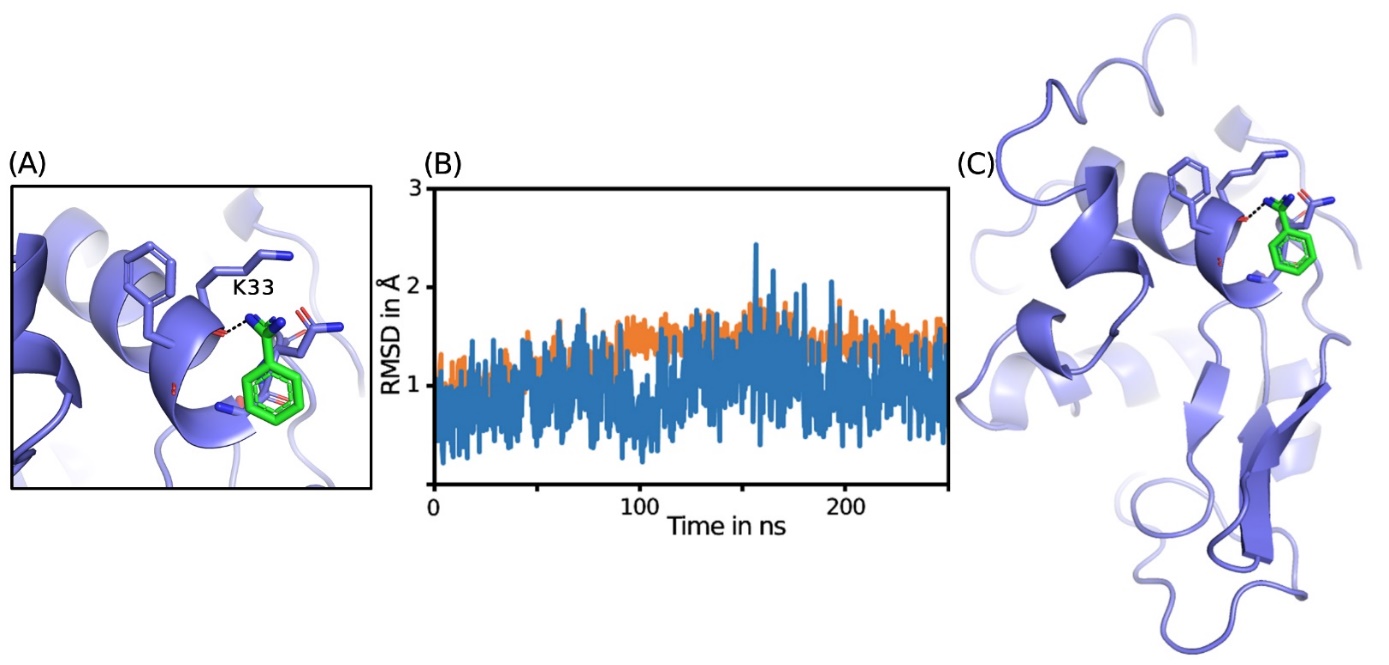

**Figure S1:**MD simulations of BEN-HEWL crystal structure. **(A)** Binding mode of BEN (green) with HEWL (slate blue), highlighting interactions with residue K33. (B) RMSD of HEWL in complex with BEN over a period of 250 ns. **(C)** Cartoon representation of the binding pose of BEN within the HEWL structure, illustrating the overall conformation and orientation of the ligand relative to the protein.
